# Collectin-11 regulates osteoclastogenesis and bone maintenance via a complement-dependent mechanism

**DOI:** 10.1101/2025.05.08.652605

**Authors:** Mark C Howard, Conrad A Farrar, Christopher L Nauser, Yusun Jeon, Anastasia Polycarpou, Dorota Smolarek, Roseanna Greenlaw, Peter Garred, Daniela A Vizitiu, Subhankar Mukhopadhyay, Steven H Sacks

## Abstract

The human developmental disorder 3MC syndrome is characterized by skeletal deformities associated with a deficiency of the pattern recognition molecule collectin-11 (CL-11), yet the underlying molecular and cellular mechanisms remain unclear. Here, we demonstrate that CL-11 deletion alone does not cause bone abnormalities in mice; however, combined deficiencies involving CL-11 and complement components MASP-2 (lectin pathway), CFB, or C3 (alternative amplification pathway) lead to significant vertebral bone loss and spinal curvature by 12 weeks of age. Ex vivo osteoclast (OCL) differentiation from bone marrow-derived cells of these double-knockout (DKO) mice was markedly impaired, but differentiation capacity was substantially restored by supplementation with CL-11. Furthermore, CL-11 and membrane attack complex (C5b-9) deposition were co-localized to OCLs and their precursors in normal bone tissues from embryonic stages to adulthood. These findings identify CL-11 as a critical osteoclastogenesis and bone maintenance regulator in conjunction with complement system-mediated signalling pathways and highlight CL-11 as a potential therapeutic target in diseases involving dysregulated osteoclast function and bone remodelling.

**Significance statement:** Our research in mice provides new insights into how mutations in the immune surveillance molecule collectin-11 contribute to skeletal abnormalities in humans. Evidence from our study suggests that normal osteoclasts interact with collectin-11 and complement, and that the disruption of this cooperation results in impaired bone maintenance in adulthood. These findings not only advance our understanding of osteoclast function but also highlight the therapeutic potential of targeting collectin-11 in conditions associated with osteoclast dysfunction.

## Introduction

Collectin-11 (CL-11) is a soluble C-type lectin that recognizes carbohydrate motifs on both self and nonself structures, activating the lectin complement pathway (LP) (Keshi et al., 2006). It plays key roles in innate immunity, inflammatory disorders, and embryogenesis (Hansen et al., 2016). The widespread tissue expression of CL-11 suggests its fundamental biological importance across different cell populations (Nauser and Sacks, 2023).

The basic structural unit of CL-11 is a triple-chain complex, with each chain comprising a carbohydrate-recognition domain (CRD) and a collagen-like domain (CLD), separated by a neck region. These CL-11 complexes self-combine into dimers and trimers (100–200 kDa), enhancing carbohydrate-binding avidity. Serine proteases (MASPs 1–3) physically associate with the CLD of CL-11, facilitating the formation of the classical pathway C3 convertase (C4bC2b), which leads to C3 cleavage (Hansen et al., 2010, Ma et al., 2013). MASP-2, besides cleaving C4 and C2, can directly cleave C3 (Ma et al., 2013, Schwaeble et al., 2011, Asgari et al., 2014). MASP-3 also contributes to the alternative pathway (AP) by activating factor D, which converts C3bB into C3bBb, the alternative pathway C3 convertase (Takahashi et al., 2010). Thus, CL-11/MASP complexes can initiate both classical and alternative C3 convertases at the ligand-binding site for CL-11 (Supplemental Figure 1).

The subsequent cleavage of C3 and C5 generates anaphylatoxins (C3a and C5a), the opsonin C3b, and membrane attack complex (C5b-9, MAC), promoting cell activation, damage, and cell death. Given that CL-11 is expressed in cells of ectodermal, endodermal, and mesenchymal origin (Hansen et al., 2016, Keshi et al., 2006), this locally produced pattern-recognition molecule (PRM) may exert regional functions. Notably, mice lacking tissue CL-11 resist post-ischemic kidney injury (Farrar et al., 2016). In humans, mutations in *COLEC11* (or the related *COLEC10* gene) cause 3MC syndrome (Rooryck et al., 2011, Atik et al., 2015, Munye et al., 2017), a developmental disorder characterized by craniotiacial dysmorphia, cleft palate, stunted growth, and multiorgan defects (Urquhart et al., 2016). The *MASP1* gene encodes three splice variants, MASP-1, MASP-3 and MAP-1. Mutations in *MASP1* making MASP-3 inactive have also been identitiied in 3MC patients (Gardner et al., 2017, Rooryck et al., 2011), suggesting that disruption of complement activation—rather than a direct effect of PRMs on developing tissues—may underlie the syndrome’s pathology.

A role for complement in bone formation is further supported by evidence that complement factors B (CFB), C3, C5, and C9 are expressed in the growth plate, a specialized cartilaginous region at the ends of long bones essential for longitudinal growth and endochondral ossification (Andrades et al., 1996, Mödinger et al., 2018). Chondrocytes also express complement components C4, C2, and C3 (Bradley et al., 1996). Moreover, mice lacking C3 or C5 exhibit impaired fracture healing, with C5 deficiency having a greater impact than C3 deficiency, highlighting the role of complement activation—particularly its terminal effector pathway—in bone remodelling (Ehrnthaller et al., 2013). Osteoclast (OCL) formation may also be directly influenced by anaphylatoxins, even in the absence of key differentiation factors RANKL and M-CSF (Ignatius et al., 2011).

To further investigate the interplay between the LP and AP triggered by CL-11, we generated double knockout (DKO) mice with disruptions in *Colec11* and either *Masp-2* (LP), *Cfb* (AP), or *C3* (common to both pathways). A striking proportion of the adult DKOs developed spinal curvature (kyphoscoliosis), whereas most single knockout (SKO) mice lacking CL-11, MASP-2, CFB, or C3 in isolation displayed no such skeletal phenotype. The present study determined how, when, and where CL-11 contributes to this unexpected skeletal phenotype.

## Results

### DKO Mice Lacking CL-11 Exhibit Loss of Vertebral Bone Integrity

We generated four groups of DKO mice: *Colec11*^*-/-*^*/C3*^*-/-*^; *Colec11*^*-/-*^*/Masp-2*^*-/-*^; *Colec11*^*-/-*^*/Cfb*^*-/-*^; *C3*^*-/-*^*/Masp-2*^*-/-*^). By 12 weeks of age, kyphosis and/or scoliosis occurred in 23–28% of the three DKO groups lacking CL-11 (*Colec11*^*-/-*^*/C3*^*-/-*^; *Colec11*^*-/-*^*/Masp-2*^*-/-*^; *Colec11*^*-/-*^*/Cfb*^*-/-*^), compared to only 0–1% in wild-type (WT) mice, SKO mice, and DKO mice with intact CL-11 (*C3*^*-/-*^*/Masp-2*^*-/-*^) (Table 1). Gross dissection and skeletal staining confirmed the spinal deformity, revealing a marked loss of distinction between bone and cartilage (Figure 1a, 1b). Thus, the absence of CL-11 predisposed mice to the skeletal phenotype, but only in combination with a second deletion (C3, CFB, or MASP-2).

**Figure 1.**
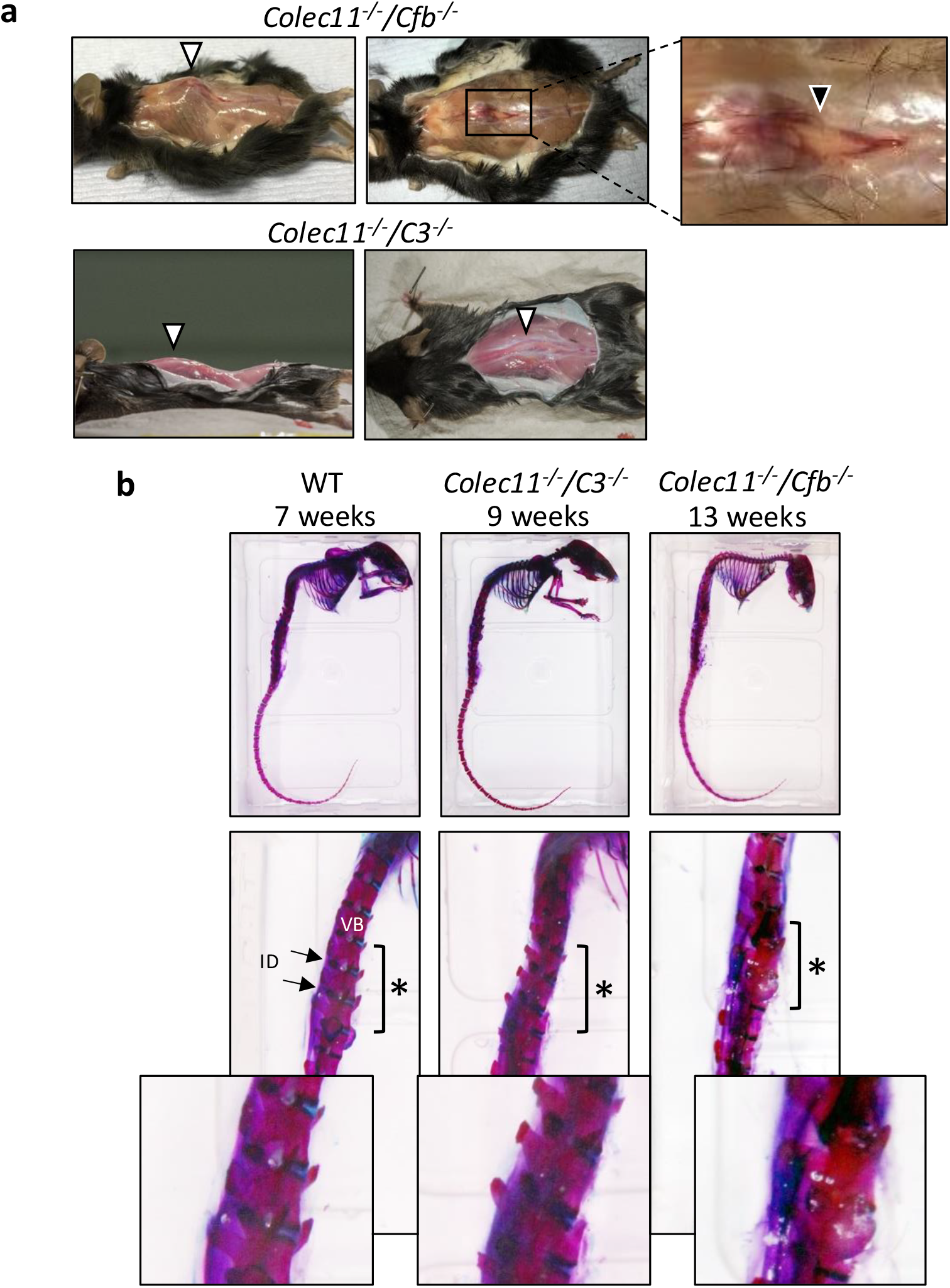
Phenotype of mice with single and double deletions of complement genes. (A) Representative images of the spinal phenotype in *Colec11*^*-/-*^/*Cfb*^*-/*-^and *Colec11*^*-/-*^/*C3*^*-/-*^and double-knockout (DKO) mice, showing kyphosis (upper panel) and scoliosis (lower panel). Lesions are exposed through skin incisions (n = 3 mice per group). Arrows indicate regions of enhanced spinal curvature. Magnified insert highlights increased fat deposits (black arrow). (B) Skeletal preparations of representative mice at different ages. Vertebral bodies (VBs) are stained with Alizarin Red, and intervertebral discs (IDs) with Alcian Blue (n = 5–7 mice per group). Stars indicate regions where VBs have distinct outlines and IDs are separate from VBs in WT and young DKO mice. In older DKO mice, VBs lose defined edges and blend with adjacent IDs. Magnified inserts highlight these regions.

Whole-body μCT imaging further highlighted the skeletal abnormalities, with vertebral bodies most affected. These vertebral bodies appeared punctuated by numerous small lesions ranging from isolated defects to extensive structural disruption (Figure 2a, 2c). Spinal processes also displayed destructive lesions. Multi-angle views allowed quantification of affected vertebrae (Figure 2b), with the lumbar and thoracic regions most severely impacted (Figure 2d). In contrast, scapular and pelvic bones exhibited only minor defects, while femurs and skull calvaria appeared unaffected (Supplemental Figure 2). Femur length did not significantly differ between KO and WT mice of the same sex.

**Figure 2.**
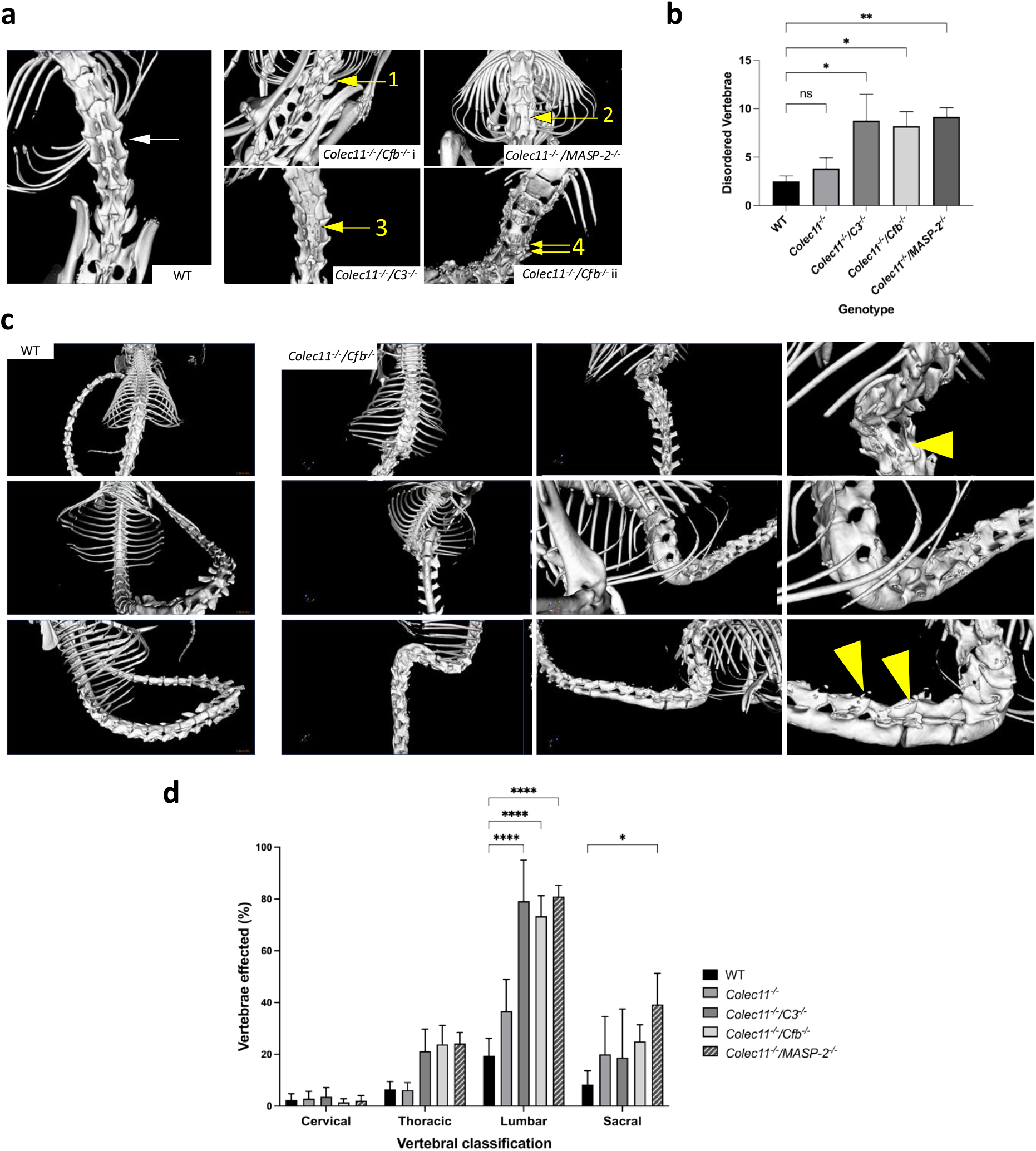
Micro-computed tomography (micro-CT) scans of affected and unaffected mice. (A) Vertebrae in WT and DKO mice, with severity ranging from mild abnormality to severe spinal disorganization. Arrow 1: normal vertebra; Arrow 2: small hole and deformed transverse processes; Arrow 3: “peppering” from multiple small holes; Arrow 4: complete vertebral breakdown. (B) Number of vertebrae affected by structural abnormalities (n = 6–11 mice per group). (C) Multiple views of the spine in a *Colec11*^*-/-*^*/Cfb*^*-/-*^mouse, showing vertebral and spinous process damage (yellow arrowheads), with the anterior surface of the vertebrae spared. (D) Distribution of vertebral regions affected by abnormalities (as in 2A, C). Each region is represented as a percentage of vertebrae containing structural abnormalities. Error bars = SEM. *p < 0.05, **p < 0.01, ****p < 0.0001.

Macroscopic and histological examinations of major organs revealed no evidence of a systemic disorder (Supplemental Figure 3). However, litter sizes were reduced by approximately one pup in DKOs lacking CL-11 compared to WT, SKO, and *C3*^*-/-*^*/Masp-2*^*-/-*^mice, despite shared housing and genetic background (Supplemental Figure 4a). Although birth weight and growth rates were lower across all KO strains compared to WT, sex ratios remained balanced (Supplemental Figure 4). Notably, skeletal abnormalities did not worsen with aging beyond 12 weeks, as confirmed by gross dissection, skeletal preparation, and μCT imaging of 12-month-old mice (n = 5/group) (data not shown).

### Vertebral Growth Plates in DKO Mice Exhibit Trabecular Bone Loss

Histological analysis of affected *Colec11*^*-/-*^DKO spines revealed significant trabecular bone loss in vertebral bodies (VB), accompanied by regions of increased marrow adiposity (Figure 3a-d). Intervertebral discs (ID), visualized with toluidine blue staining, showed degeneration in the nucleus pulposus (NP). Thus, vertebral trabecular bone thinning, replacement by adipose tissue, and ID degeneration characterize the destructive spinal lesions in *Colec11*^*-/-*^DKOs.

**Figure 3.**
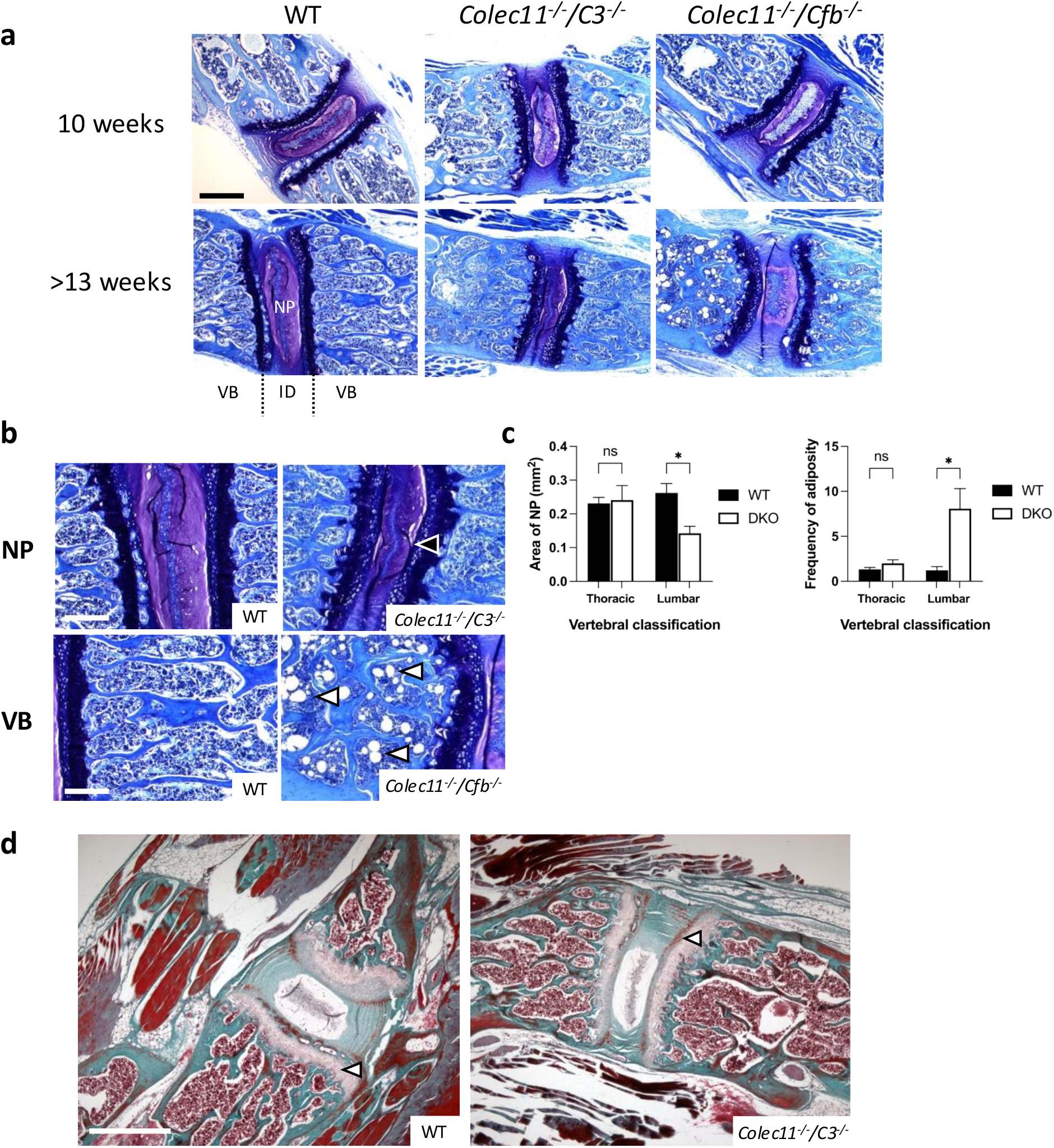
Histological characterization of bone phenotype and intervertebral disc lesions in affected mice. (A, B) Longitudinal sections of vertebral bodies (VB) and intervertebral discs (ID) stained with toluidine blue. (A) Images of WT and DKO mice at 10 weeks (upper panel) and 14–20 weeks (lower panel) (scale bar = 500 μm). (B) Higher magnification images from (A) of older mice, showing nucleus pulposus (NP) degradation (black arrow), loss of structural organization, and increased adiposity in DKO mice (white arrows). (C) Quantification of NP degradation and increased VB adiposity. (D) Masson-Goldner staining of VBs, showing similar osteoid thickness (white arrows) (n = 3–5 mice per group, scale bar = 250 μm). Error bars = SEM. *p < 0.05.

### Embryonic CL-11 and C3 Deficiency Do Not Affect Bone Morphology

To determine whether these defects originated during embryonic development, we examined embryonic tissues from *Colec11*^*-/-*^*/C3*^*-/-*^mice. Light microscopy revealed no overt morphological abnormalities, despite CL-11 and C3 being broadly expressed in WT embryonic vertebrae (Figure 4a, 4b). Between E13.5–E18.5, CL-11 and C3 staining was prominent in regions of condensing chondrocytes, later becoming restricted to primary ossification centres (Figure 4b, 4c).

**Figure 4.**
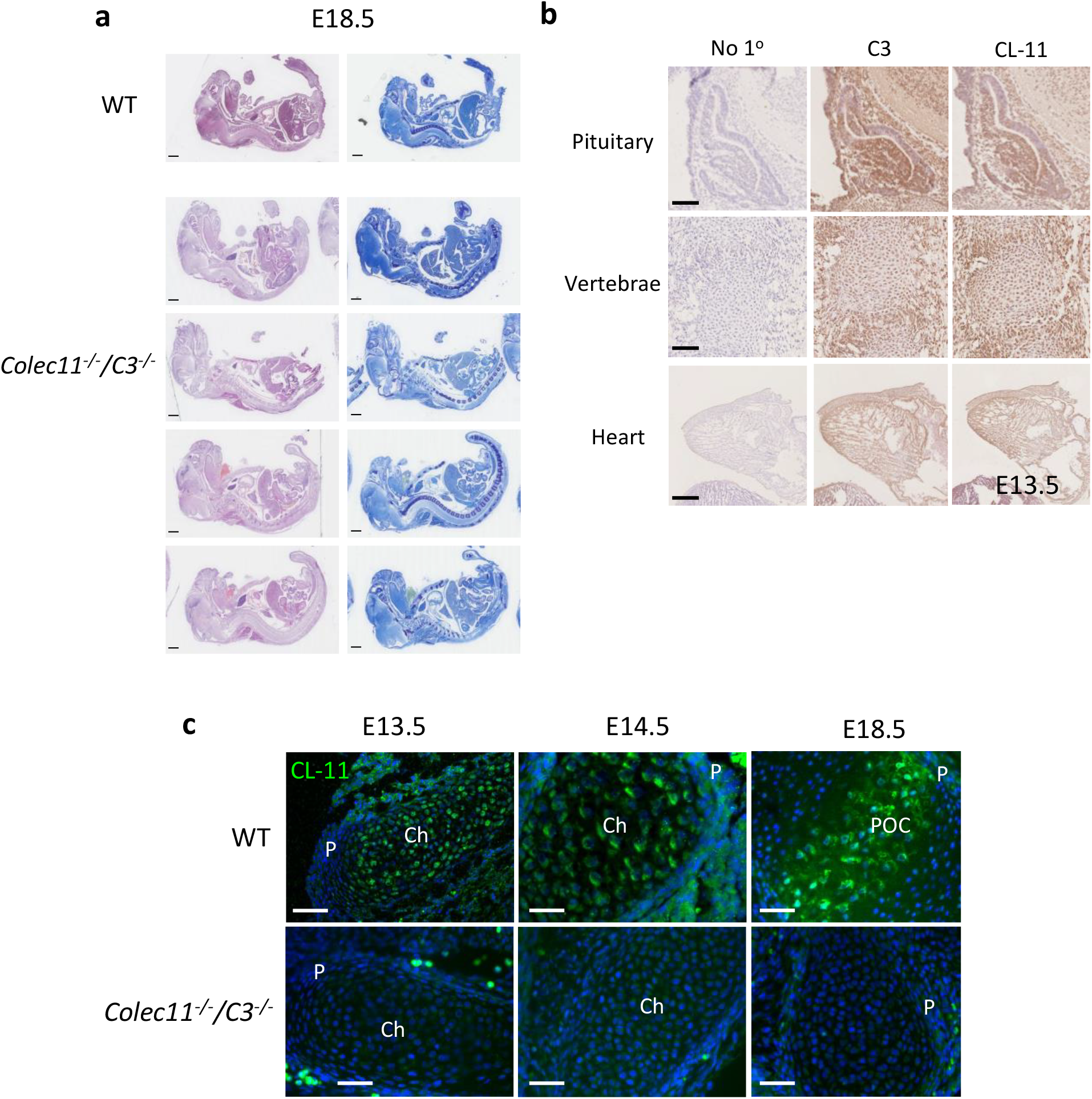
Phenotype and complement staining for CL-11 and C3 in developing embryos. (A) Sagittal sections of E18.5 WT and DKO embryos stained with H&E and toluidine blue, showing no overt embryonic phenotype (scale bar = 1000 μm). (B) Immunostaining for C3 and CL-11 in specific E13.5 WT tissues, including pituitary, heart and presumptive spinal column (scale bar = 100 μm). “No 1°” indicates control without primary antibody. (C) CL-11 staining of long bones at E13.5, E14.5, and E18.5. CL-11 localizes to chondrocytes (Ch) at E13.5 and E14.5, appears in the perichondrium (P) at E14.5, and concentrates in the primary ossification centre (POC) at E18.5. *Colec11*^*-/-*^*/C3*^*-/-*^tissues serve as antibody controls, showing autofluorescence in blood cells but no specific staining in bone tissue (scale bar = 100 μm).

To assess MAC formation, we probed embryonic tissue sections with a C9-specific antibody. C9 aggregates were detected dynamically between E13.5–E18.5 (Figure 5a). WT tissues showed large pericellular C9 aggregates indicative of MAC deposition, with progressive regional restriction over time, leading to an overall reduction in MAC area (Figure 5b). In contrast, *Colec11*^*-/-*^*/C3*^*-/-*^embryos exhibited markedly reduced C9 staining at all stages. However, SKO *Colec11*^*-/-*^embryos did not show a significant decline in C9 over time. Thus, MAC formation was impaired only in the absence of both CL-11 and C3—but without evident morphological consequences by E18.5.

**Figure 5.**
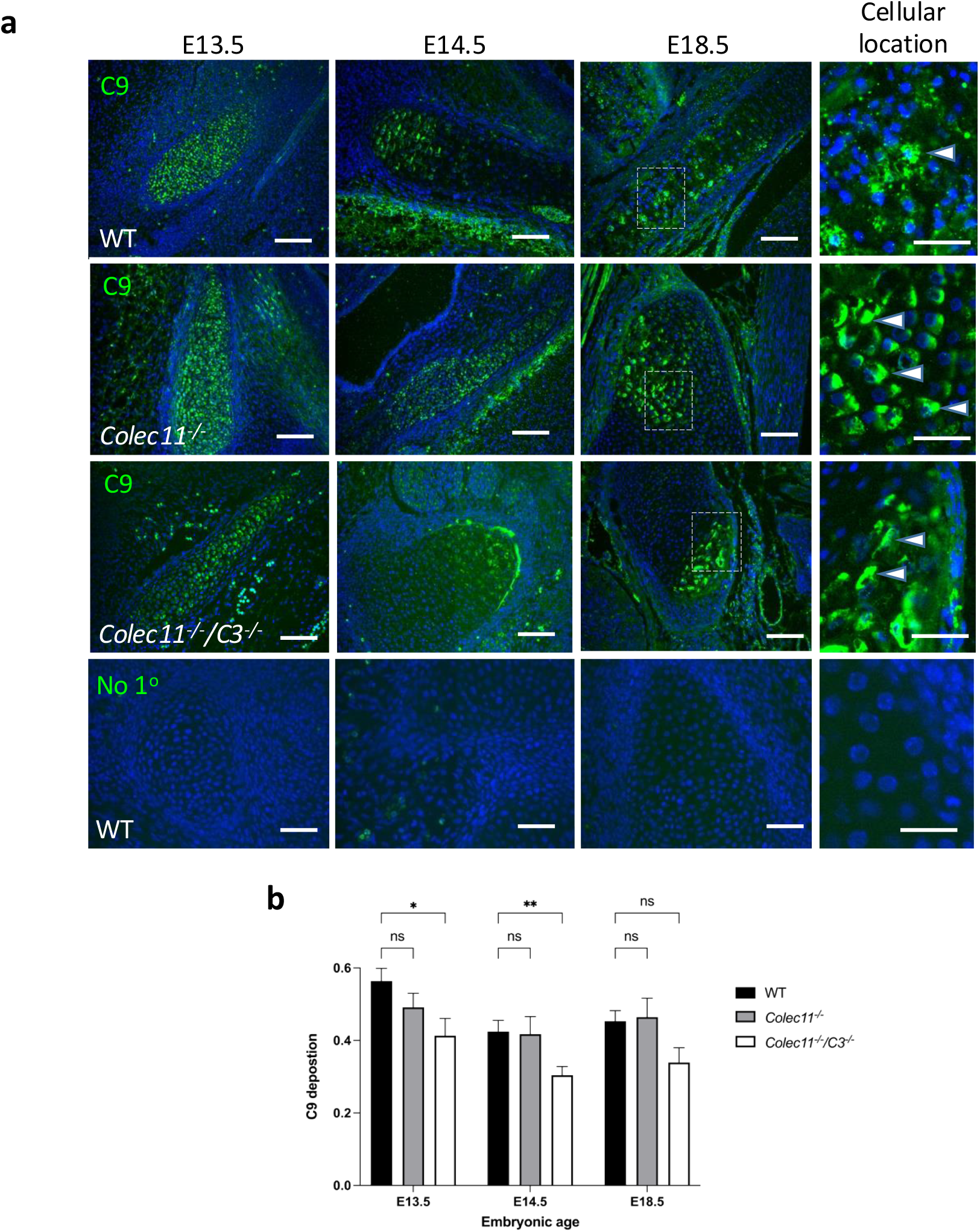
Membrane Attack Complex (MAC) detection in embryonic bone. (A, B) C9 staining in long bones at different embryonic stages (scale bar = 100 μm). Clumped C9 staining (white arrows) suggests polymeric C9 incorporation into MAC (scale bar = 50 μm). *Colec11*^*-/-*^*/C3*^*-/-*^mice show significantly reduced C9 deposition at E13.5 and E14.5, while *Colec11*^*-/-*^mice show no major reduction (n = 3–4 mice per group). The last row shows WT embryos stained without primary antibody to control for secondary antibody specificity. Error bars = SEM. *p < 0.05, **p < 0.01.

### Osteoclast Differentiation Requires CL-11 and Either C3 or CfB

Since skeletal abnormalities emerged postnatally (≥ 12 weeks), we investigated a potential defect in bone maintenance rather than development. Given prior links between complement and OCL formation (Ignatius et al., 2011), we assessed bone marrow-derived stem cell (BMSC) differentiation into OCLs across genotypes. WT BMSCs differentiated into mature multinucleated giant cell OCLs, with concurrent CL-11 expression, OSCAR detection, and TRAP^+^ staining (Figure 6a–c). However, in *Colec11*^*-/-*^*/C3*^*-/-*^BMSCs the number of multinucleated OCLs was significantly reduced as indicated by the microscopic area occupied by such cells (Figure 6d and 6e). MAC formation, detected via C9 staining, was also impaired in *Colec11*^*-/-*^*/C3*^*-/-*^myeloid progeny (Figure 6f, 6g). Similarly, *Colec11*^*-/-*^*/Cfb*^*-/-*^cells largely failed to differentiate into mature OCLs (Figure 6e; Supplemental Figure 5a). In contrast, OCLs derived from *Colec11*^*-/-*^SKOs were only mildly reduced in number and MAC deposition (Figure 6d–g; Supplemental Figure 5b). These findings suggest that OCL differentiation is dependent on CL-11 facilitated by either C3 or CFB.

**Figure 6.**
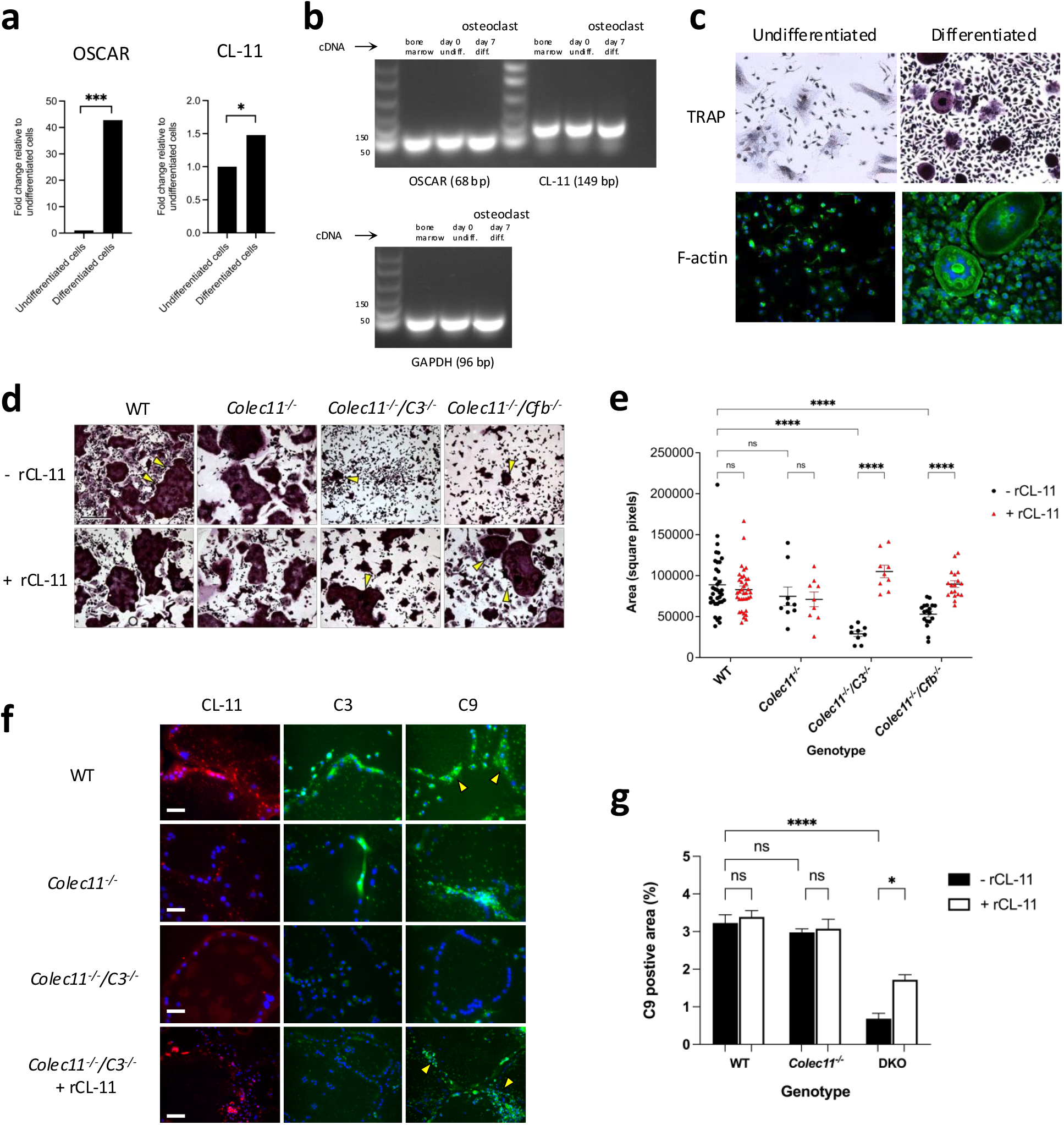
Osteoclast (OCL) differentiation from adult bone marrow stem cells and CL-11 impact. (A) qPCR analysis of OSCAR and CL-11 expression in undifferentiated and differentiated WT cells. Bars represent means from two technical replicates per gene (n = 12 coverslips). Data is representative of two independent experiments. (B) Agarose gels of qPCR products with GAPDH as a loading control. (C) Microscopy of cells stained for TRAP and F-actin (scale bar = 100 μm). (D) Representative images of OCL differentiation in WT, *Colec11*^*-/-*^, *Colec11*^*-/-*^*/C3*^*-/-*^and *Colec11*^*-/-*^ */Cfb*^*-/-*^cells ± rCL-11 addition. Arrows indicate multinucleated OCLs with size changes in DKO cultures (scale bar = 100 μm). Each row of images in each treatment group represents a single mouse. (E) Quantification of OCL area in square pixels. Each dot represents one image (n=6 WT, n=3 *Colec11*^*-/-*^, n=1 *Colec11*^*-/-*^*/C3*^*-/-*^and n=2 *Colec11*^*-/-*^*/Cfb*^*-/-*^mouse). Data were analysed using two-way ANOVA with Sidak’s multiple comparisons. (F) Immunostaining of cultures for CL-11, C3, or C9, showing C9 aggregates on OCLs (scale bar = 250 μm). (G) Quantification of C9 staining in (F). Error bars = SEM. *p < 0.05, ***p < 0.001, ****p < 0.0001.

### Exogenous CL-11 Restores Osteoclast Differentiation

To determine whether CL-11 directly supports OCL differentiation, we supplemented the differentiation medium with recombinant CL-11 (rCL-11, 0.9 μg/mL) in *Colec11*^-/-^/*C3*^-/-^ and *Colec11*^-/-^/*Cfb*^-/-^ BMSC cultures. This treatment partially restored CL-11 binding and significantly increased the presence of multinucleated giant cell OCLs (Figure 6d and 6e; Supplemental Figure 5). Consistent with this observation, the average number of nuclei per OCL in rCL-11–treated *Colec11*^-/-^/*C3*^-/-^ BMSCs increased to 12.6, compared with 6.6 in untreated *Colec11*^-/-^/*C3*^-/-^ cells (p < 0.01) and 16 in WT cells (p < 0.001 vs. untreated *Colec11*^-/-^/*C3*^-/-^ cells). Cell-surface membrane attack complex (MAC) deposition was also partially rescued (Figure 6f, 6g). Notably, since rCL-11 restored MAC formation even in the absence of added C3 (Figure 6f), this suggests an alternative, C3-bypass activation mechanism downstream of CL-11 that leads to activation of the terminal complement pathway.

### CL-11 and Complement Interact with Osteoclasts Throughout Life

Given these ex vivo findings, we hypothesized that CL-11–OCL interactions could activate complement and promote OCL differentiation under physiological conditions. To test this, we analysed unmanipulated WT mouse bone sections for CL-11 and MAC deposition. Large multinucleated TRAP^+^ OCLs were strongly positive for CL-11 and polymeric C9 (Figure 7a). Similarly, in normal embryonic bone at different pre-OCL developmental stages, CL-11 and C9 localized to regions of cellular condensation within presumptive vertebral bodies (Figure 7b).

**Figure 7.**
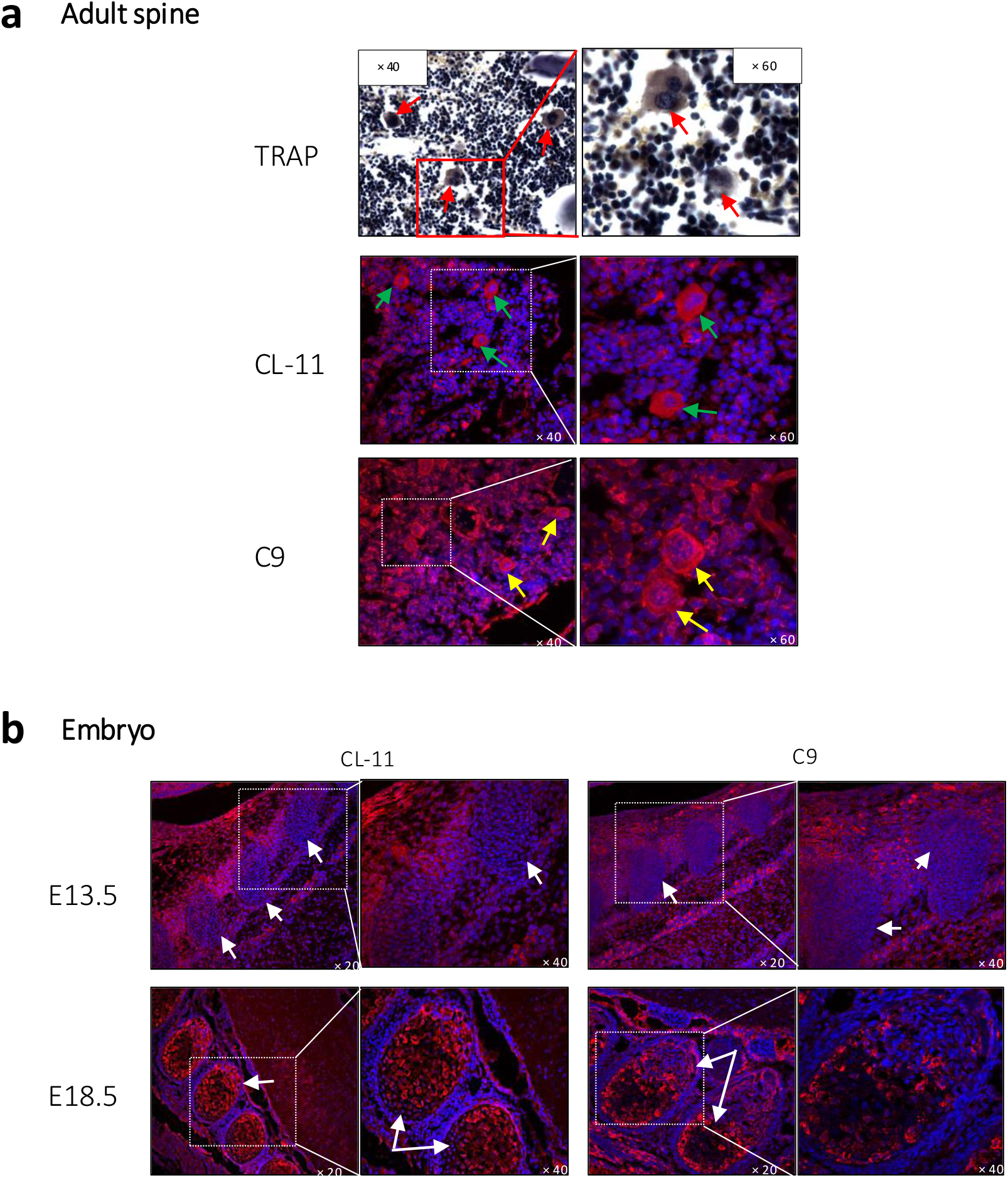
Localization of osteoclasts, CL-11, and MAC in adult and embryonic spines. (A) TRAP-stained adult vertebral bodies, showing multinucleated OCLs (red arrows), CL-11 (green arrows), and C9 (yellow arrows) (n = 3). (B) Confocal images of embryonic vertebral bodies at E13.5 and E18.5 (white arrows). Adult spines contain differentiated OCLs with CL-11 and MAC deposition. By E18.5, vertebral condensation begins, with CL-11 and MAC-positive progenitor cells present.

These findings strongly support a functional relationship between CL-11, MAC assembly, and OCL differentiation. Our data suggest that CL-11 and complement play a regulatory role in osteoclastogenesis from embryonic bone development through adult bone maintenance. We propose that chronic failure of this function leads to vertebral pathology by 12–13 weeks of age, as observed in our mouse model.

## Discussion

Our results demonstrate that mice lacking CL-11 develop age-related vertebral bone loss, but only in the absence of at least one additional key component of the lectin or alternative complement pathways. This dual deficiency was associated with impaired osteoclast (OCL) differentiation in bone marrow cultures, which was largely restored by supplementation with recombinant CL-11 at physiological concentration. The presence of CL-11 and the membrane attack complex (C5b-9) on osteoclasts and precursor cells in normal bone further supports the role of CL-11 as a physiological regulator of OCL differentiation, contributing to bone remodelling throughout life.

Not all mice with the vulnerable double-knockout (DKO) genotype were affected, likely due to genetic and environmental interplay leading to overt bone pathology in up to 28% of adults. The lumbo-thoracic regions of the vertebral spine were most susceptible, suggesting that repetitive mechanical stress from body movements during feeding and grooming may contribute to the phenotype.

Bone is a dynamic tissue maintained by bone-forming osteoblasts (OBLs) and bone-resorbing OCLs, both of which interact with the complement system. OBLs, derived from mesenchymal lineage, secrete bone matrix before differentiating into osteocytes (Schoengraf et al., 2013). OBLs are potential targets for C5a, as C5aR1 is involved in osteogenic differentiation and migration (Bergdolt et al., 2017, Ignatius et al., 2011). In contrast, OCLs arise from the monocyte-macrophage lineage and they degrade bone matrix through proteolytic enzymes and acidification (Schoengraf et al., 2013). Deletion of CD59, an inhibitor of MAC formation, results in increased bone resorption and cortical bone excess due to elevated OCL differentiation, with no impact on OBL differentiation (Bloom et al., 2016). Our data suggest that the complement system plays a homeostatic role in bone maintenance, implicating CL-11 as a critical trigger in this mechanism. While MAC gain-of-function appears to promote excessive long bone growth (Bloom et al., 2016), our study shows that its loss leads to vertebral spine disintegration. Although both gain and loss of MAC affect the skeletal system, CL-11 has multifunctional roles beyond MAC signalling, explaining the differing bone defects.

The bone abnormalities observed in our study required the loss of a second complement activation component alongside CL-11. Redundancy is a hallmark of the complement system, ensuring effective conversion of C3 and C5 into their active forms through multiple pathways. This complexity may explain why an additional activation-pathway defect (e.g., CFB, MASP-2, C3) was necessary for the bone repair mechanism to fail. Conversely, MAC formation was restored by CL-11 supplementation, even in the absence of C3. Like thrombin—a protease capable of cleaving C5 to bypass C3 (Huber-Lang et al., 2006)—CL-11-driven LP activation may directly cleave C5, initiating MAC formation independently of C3, albeit less efficiently.

The mouse phenotype in our model is more specific than that of most of the reported 3MC syndrome patients, who present with multiple developmental defects. Besides common features such as hypertelorism and craniosynostosis (Munye et al., 2017) skeletal abnormalities in 3MC include malformed ears (Talenti et al., 2018), hypoplastic scapulae (Gardner et al., 2017), skull asymmetry (Basdemirci et al., 2019), and various cleft palate forms (Leal et al., 2008, Munye et al., 2017). Mechanistic studies have focused on neural crest cell (NCC) patterning (Nauser and Sacks, 2023, Rooryck et al., 2011), which gives rise to craniofacial mesenchyme and contributes to skull and pharyngeal bone development (Minoux and Rijli, 2010). The restricted phenotype observed in our mice, manifesting only around 12 weeks of age, suggests distinct pathogenic mechanisms beyond embryonic NCC migration. Notably, no skull abnormalities were detected in affected mice by CT measurement. Our focus on OCLs reveals a different CL-11-regulated cellular mechanism. The observed trabecular bone loss, increased marrow adiposity, and nucleus pulposus degradation of intervertebral discs align with dysregulation in both mesenchymal (e.g., OBLs, chondrocytes, adipocytes) and hematopoietic (e.g., OCLs) lineages.

Complement activation in wild-type (WT) embryos was evident between E13.5 and E18.5, with MAC formation detected on chondrocytes from E13.5—coinciding with the cartilaginous anlagen formation before primary ossification at E18.5. This period marks the transition of proliferating chondrocytes to hypertrophic chondrocytes, followed by invasion by OCLs and endothelial cells, and subsequent bone matrix deposition by differentiating OBLs (Berendsen and Olsen, 2015, Egawa et al., 2014). The reduced MAC formation observed in DKO embryos from E13.5 to E18.5 may have impaired endochondral ossification, predisposing them to mechanical stress-induced bone decay in adult life.

Bone marrow stromal cells and monocyte-macrophage lineages produce complement components (e.g., CL-11, C3, CFB) that regulate the local microenvironment. Our differentiation experiments incorporated no exogenous complement, except for rCL-11 in the reconstitution study. Given this CL-11 gene expression and detected complement activation products (C3d, MAC), we propose that complement proteins produced locally regulate OCL differentiation from myeloid precursors. Thus, OCLs and their progenitors likely regulate by secreting CL-11 and other complement factors, contributing to OCL renewal and bone maintenance.

In summary, we identify a novel function of CL-11 in regulating OCL differentiation from myeloid precursors. This function depends on synergy with other lectin or alternative pathway components; without it, bone integrity deteriorates with age. Our model offers insight into how CL-11 mutations and lack of complement signalling contribute to human skeletal abnormalities and suggest involvement of 3MC syndrome mechanisms beyond embryonic NCC migration. Conversely, excessive OCL activity is a hallmark of diseases such as renal bone disease, osteoporosis and erosive osteoarthritis, where monoclonal antibody drugs targeting the OCL differentiation factor RANKL are now in clinical use or being evaluated (Cummings et al., 2009, Wittoek et al., 2024). Our study suggests CL-11 too as a potential target in this context.

## Materials and Methods

### Animals

*Colec11*^*-/-*^mice (Farrar et al., 2016; Howard et al., 2020) were backcrossed to C57BL/6 for four generations. *Masp-2*^*-/-*^mice were kindly provided by W. Schwaeble, Cambridge, UK (Asgari et al., 2014). *C3*^*-/-*^mice were used as previously described (Zhou et al., 2000; Wessels et al., 1995). *Cfb*^*-/-*^mice were a generous gift from R. Wetsel, University of Texas (Matsumoto et al., 1997; Watanabe et al., 2000).

DKO-homozygote mice were generated by crossbreeding *Colec11*^*-/-*^mice with *C3*^*-/-*^, *Cfb*^*-/-*^and *Masp-2*^*-/-*^strains and genotyped by PCR. Kyphosis and scoliosis were identified through visual inspection and spine palpation by two independent observers in both KO and WT mice of the same age. All experiments adhered to the Animals (Scientific Procedures) Act 1986.

### Dissection and Micro-CT

A dorsal incision from the scapula to the pelvis allowed direct inspection of the spines. Computed tomography was performed using a GE Explore Locus SP μCT scanner at the KCL Craniofacial Regenerative Biology Centre (Tabler et al., 2013). Three-dimensional isosurfaces were quantified using MicroView software (GE), enabling the identification of abnormal vertebrae with interruptions in smooth surfaces. The affected scapular area was calculated as a proportion of the total area.

### Skeletal Preparations

Whole-mount skeletal staining followed the method described by Rigueur and Lyons (2014). Dehydrated skeletons were fixed in 95% ethanol overnight at room temperature (RT) twice, followed by fixation in 100% acetone at RT to remove adipose tissue. Specimens were stained with Alcian Blue (0.03% w/v in 80% ethanol and 20% glacial acetic acid) for three days, washed in 75% ethanol, destained overnight in 95% ethanol, and precleared in 1% potassium hydroxide (w/v) overnight. Alizarin Red staining (0.005% w/v; MP Biochemicals) in 1% potassium hydroxide was performed for five days. Samples were cleared in 1% potassium hydroxide overnight and stored in 100% glycerol (Sigma). Imaging was conducted using an SL1 digital camera (Canon) with specimens illuminated from behind. GNU Image Manipulation Program (GIMP) software was used to standardize light levels across specimens.

### Histology

Kidney, liver, heart, and pituitary tissues were fixed overnight in 4% paraformaldehyde (PFA) in PBS, transferred to 70% ethanol, and paraffin embedded. Bone specimens were dissected to remove soft tissue and fixed overnight in 4% PFA. Long bones were decalcified in 14% EDTA (w/v; Sigma) for 14 days (pH 7.2), with daily EDTA replacement. Samples were rinsed four times in dH_2_O, dehydrated through graded ethanol (30%-70%), and paraffin-embedded. Embryonic tissue was fixed overnight in 4% PFA and transferred to 70% ethanol before embedding.

Sections (4-7 μm) were stained with haematoxylin-eosin, toluidine blue, or Masson-Goldner stain (Sigma-Aldrich; 1.00485). Toluidine blue was used to detect proteoglycans/glycosaminoglycans in cartilage, while Masson-Goldner staining identified muscle fibres, collagenous fibres, fibrin, and erythrocytes. Haematoxylin staining was used for vertebral body visualization.

### Immunohistochemistry and Immunofluorescence

For immunohistochemistry, antigen retrieval was performed on deparaffinized tissue sections using 100 mM sodium citrate (pH 6.0), followed by transfer to PBS. Tissue sections were blocked for 1 hour before incubation with primary antibodies (Table 2) overnight at 4°C. A biotinylated secondary antibody was applied for 1 hour at RT, followed by ABC kit and DAB (Vector Labs), counterstaining with haematoxylin, and dehydration for mounting with DPX (Sigma).

For immunofluorescence, a biotinylated anti-rabbit secondary antibody (Vector Labs; BA-1000) was applied for 1 hour at RT, followed by FITC-conjugated streptavidin (GeneTex; GTX30950) or streptavidin-594 (Vector Labs; SA-5594). Nuclear staining was performed using DAPI (1:10,000 in PBS; Life Technologies) before mounting with PermaFluor (LabVision). Mouse anti-human CL-11 staining employed a mouse-on-mouse kit (Vector Labs; BMK-2202). Spinal sections were stained with TRAP (Sigma-Aldrich; 387A) for osteoclast identification.

### Bone Marrow Extraction

Bone marrow (BM) was extracted from femurs and tibias of euthanized mice following Maridas et al. (2018). Bones were cleared of soft tissue, washed in 70% ethanol for 1 minute, and transferred to PBS. BM was flushed out using a 25G needle, filtered through a 70 μm strainer, centrifuged (10 min at 1200 rpm), resuspended at 2 × 10^6^ cells/mL in BM medium (10% FBS, 1% Pen/Strep in α-MEM [Gibco]), and preincubated for 2 hours on a 13 mm glass coverslips coated with 1% gelatin (w/v) (Howard et al., 2020). Media was changed after 72 hours and subsequently every 48 hours until confluency.

### Osteoclast Differentiation

Osteoclast differentiation was induced using BM medium supplemented with 0.05 μg/mL murine RANKL (PeproTech; 315-11C) and 0.015 μg/mL murine CSF (Life Technologies; RP8615), replaced every 48 hours for seven days. Rescue experiments included 2 mM CaCl_2_ (Sigma-Aldrich) and 0.9 μg/mL human rCL-11 (Howard et al., 2020; Venkatraman Girija et al., 2015) at each medium replacement. Staining was performed for TRAP or Alexa Fluor 488-conjugated phalloidin (Thermo-Fisher Scientific; A12379) and photographs were taken using an Olympus BX51 fluorescent microscope.

### Quantitative Real-Time PCR

RNA was extracted from BM-derived cells using the miRNeasy kit (Qiagen; 217084), followed by cDNA synthesis (GoScript Reverse Transcription System, Promega; A500). Gene expression was quantified using the 2-ΔΔCt method, with GAPDH as the reference gene. Results are expressed as fold-change in expression of test genes in differentiated versus undifferentiated cells.

### Quantification of Staining

ImageJ (NIH) software (Howard et al., 2020; Farrar et al., 2016) in conjunction with Ilastik 1.4.1rc2 software were used to generate pixel-based segmentation images of TRAP-stained OCLs and calculate the area of OCLs in microscopic images —see Supplemental Fig 5c for details. Percent C9 staining was normalized to DAPI. Vertebral adiposity was estimated by counting circular white areas in at least three vertebrae across six sections per vertebral region. Nucleus pulposus area was measured using NDP Viewer (Hamamatsu Photonics). Osteoclast quantification was based on TRAP-stained cultures at 100× magnification.

### Statistics

Data are presented as mean ± SEM. Comparisons between two groups were analysed using an unpaired two-tailed Student’s t-test (p < 0.05 considered significant). One-way or two-way ANOVA with multiple comparisons was used for three or more groups (p < 0.05 considered significant). Statistical analysis was conducted using GraphPad Prism v9.

## Summary of Supplemental Material

Supplemental figures include CT images of extra-vertebral bones (Supplemental Fig 2), histological sections of DKO soft tissues (Supplemental Fig 3), litter size and growth rates (Supplemental Fig 4), and differentiation study results (Supplemental Fig 5). A graphical abstract is also provided.

## Data Availability Statement

All relevant data are included in the main manuscript and/or Supplemental files.

## Acknowledgements

We thank Christopher Howard for his guidance on image analysis and Professor Paul Morgan for donating the C9 antibody used in this study. We are grateful to Professor Russel Wallis for providing the rCL-11 clone. Additionally, we appreciate the valuable advice on manuscript preparation from Professor Karen Liu, Professor Agamemnon Grigoriadis, and Dr William Barrell. This work was supported by the Medical Research Council grant MR/M012263/1: *Collectin-11 as a Trigger of the Innate Immune Response in Renal Transplantation*.

## Author Contributions

M. Howard, C. Nauser, C. Farrar, S. Mukhopadhyay, and S. Sacks conceived and designed the experiments. M. Howard, A. Polycarpou, D. Smolarek, R. Greenlaw, C. Nauser, Y. Jeon, D. Vizitiu, and C. Farrar performed the experiments. M. Howard conducted the statistical analysis. Peter Garred provided a specific antibody and contributed to manuscript preparation. C. Farrar and S. Sacks assisted with result interpretation. M. Howard led the manuscript writing, with substantial input from S. Sacks. All authors provided critical feedback and contributed to shaping the research, analysis, and manuscript.

