## Supplemental figures and tables for "Collectin-11 regulates osteoclastogenesis and bone maintenance via a complement-dependent mechanism"

### Supplemental Figure Legends

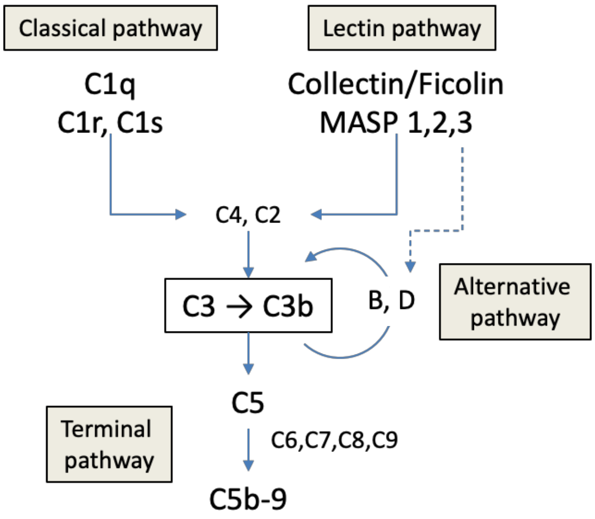


**Supplemental Figure 1. Complement activation pathways.**
In the lectin pathway of complement activation, collectin-11 (CL-11) binds carbohydrate ligands on activating surfaces. The associated serine proteases (MASPs 1-3) then facilitate the cleavage of C4 and C2, leading to the formation of C4bC2b—the C3 convertase shared with the classical pathway. The cleavage of C3 produces C3b, which contributes to the formation of C4bC2bC3b, the shared C5 convertase responsible for initiating C5b-9 membrane pore formation on the activating surface. Additionally, C3b acts as an acceptor for factor B, which, upon cleavage by factor D, forms C3bBb—the alternative pathway C3 convertase responsible for further C3b generation. Crucially, MASP-3 plays a potential role in this amplification loop by directly activating factor D.

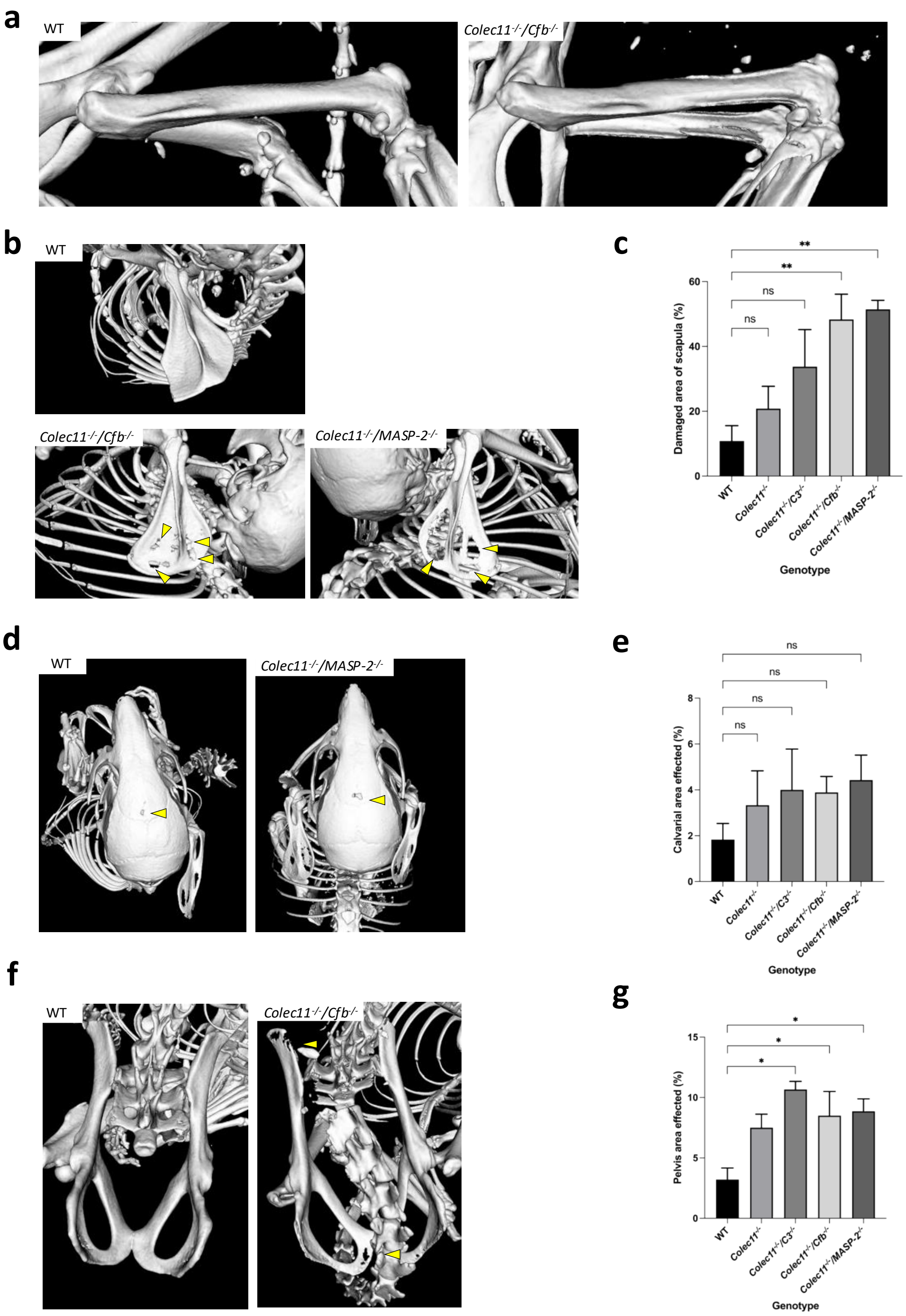


**Supplemental Figure 2. Complement deficiency affects multiple bone structures.**
Micro-CT images of femurs (A), scapulae (B, quantified in C), calvaria (D, quantified in E), and pelvis (F, quantified in G). Yellow arrowheads indicate scapular and pelvic lesions. n = 6–11 mice per group. Error bars = SEM. *p < 0.05, **p < 0.01.

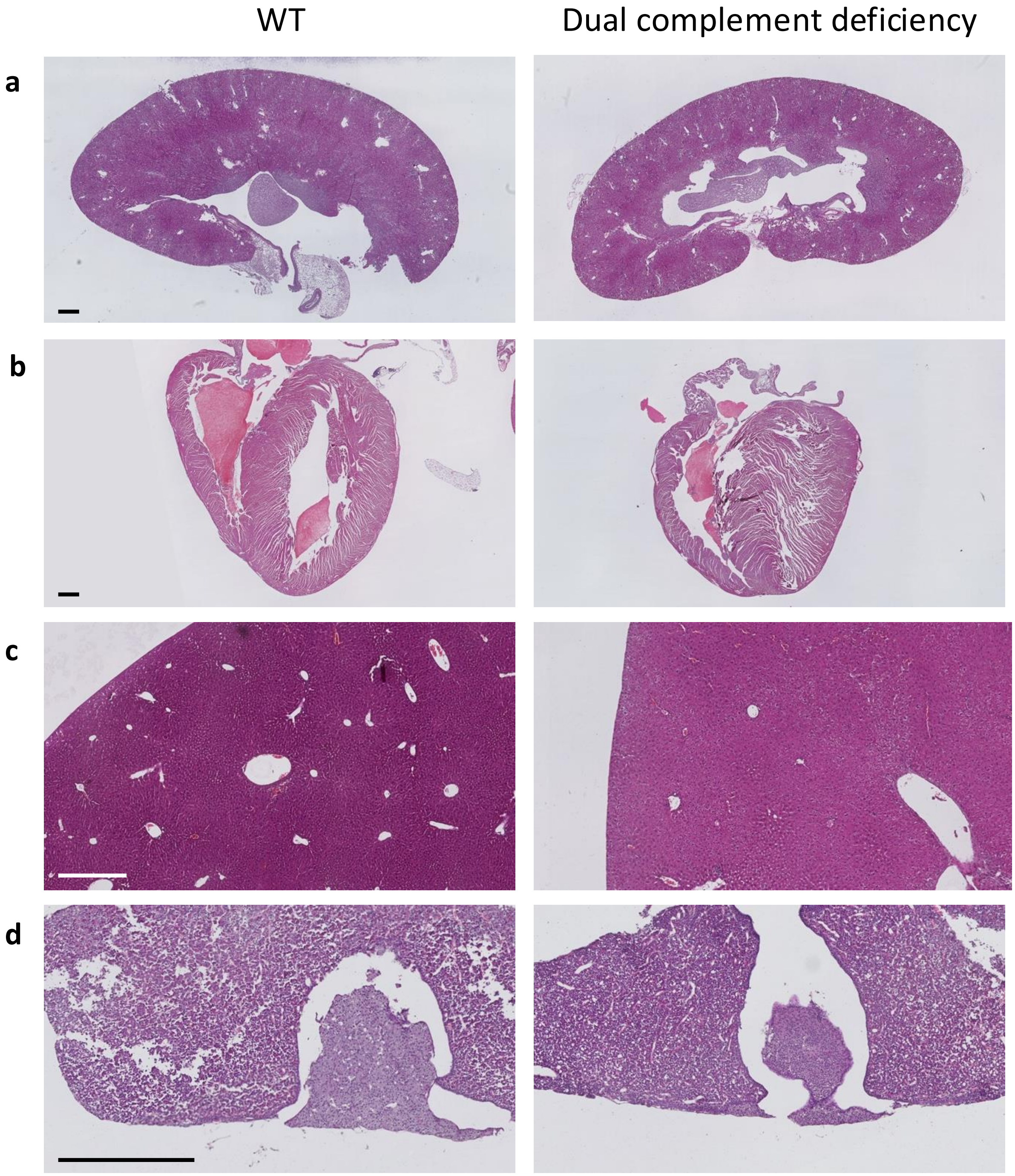


**Supplemental Figure 3. No major organ abnormalities in complement-deficient mice.** Representative images of kidney (A), heart (B), liver (C), and pituitary (D) from WT and CL-11-DKO mice (n = 6 per group, ≥12 weeks old, scale bar = 500 µm).

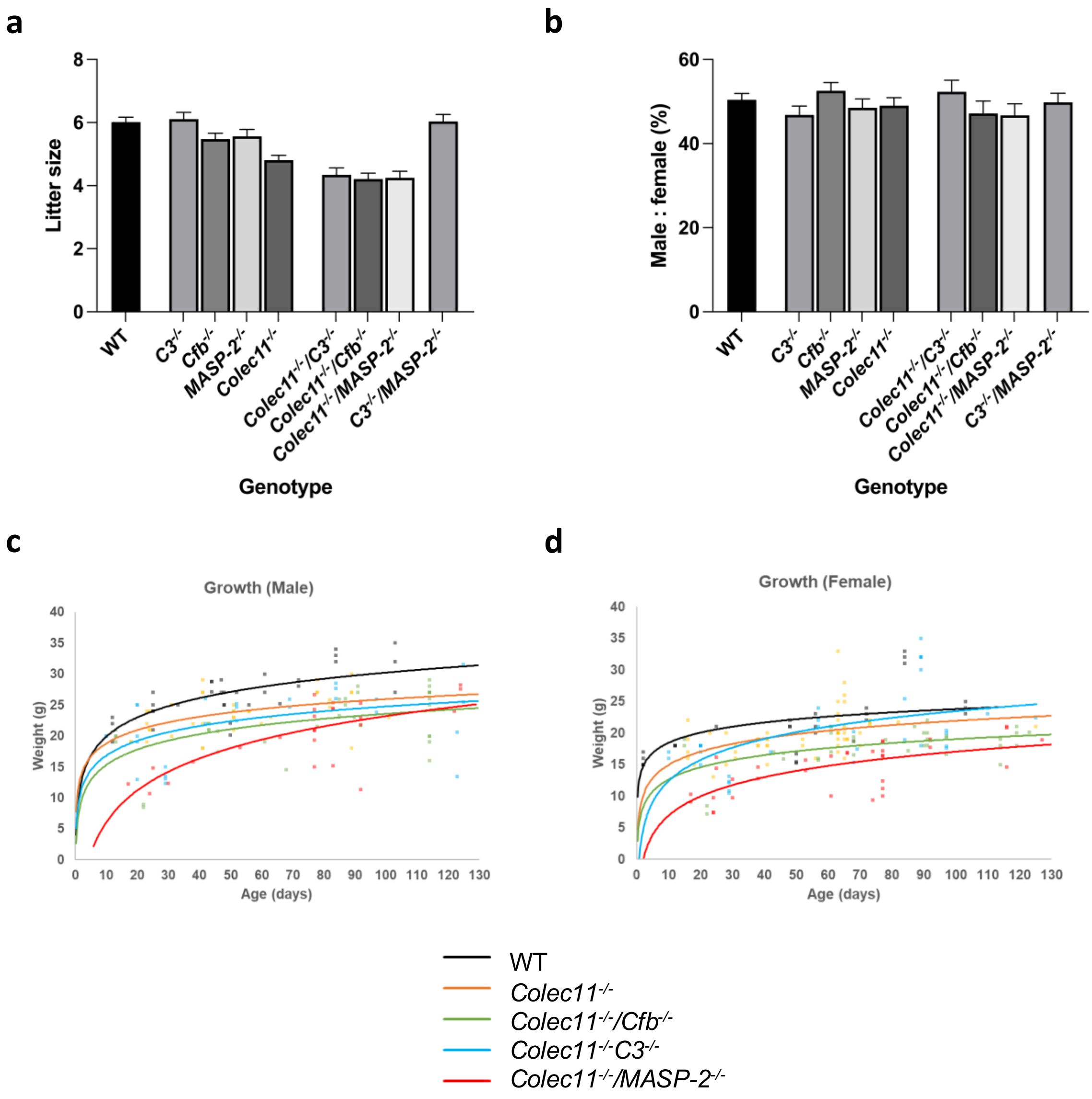


**Supplemental Figure 4. Litter size and growth rates in knockout mice.**
(A) Viable litter size by genotype. DKO mice lacking CL-11 have smaller litters.
(B) Male/female ratios indicate no sex differences (n = 77–147 litters; 319–872 mice per group).
(C, D) Growth rates in males (C) and females (D). DKO and *Colec11*$^{-/-}$ mice show lower birth rates and slower growth. Other knockouts grow similarly to WT. Error bars = SEM.

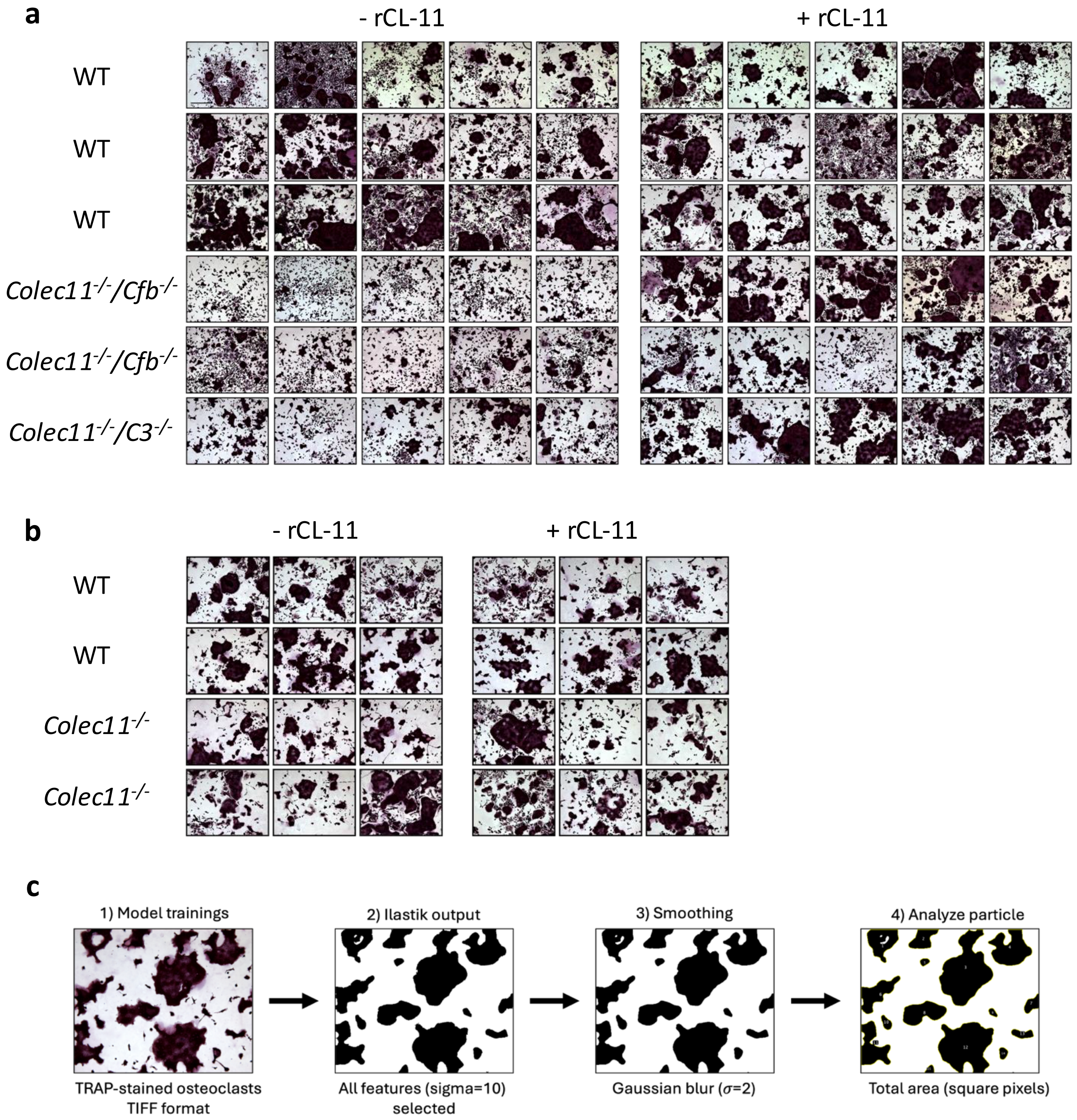


**Supplemental Figure 5. CL-11 impact on osteoclast (OCL) differentiation.**
Additional representative images of (A, B) TRAP staining of OCLs differentiated from WT and knockout BMSC at day 7 (100 × magnification, scale bar = 100μm). DKO marrow differentiates poorly without rCL-11. *Colec11*$^{-/-}$ marrow shows mildly reduced OCL differentiation. SKO marrow lacking CL-11 differentiates well even without CL-11 addition. Each row represents a single mouse. (C). Method for calculating area of giant cell OCLs presented in Fig 6e of the manuscript.
Pixel classification using ilastik (version 1.4.1rc2) was employed to generate automated segmentation images of TRAP-stained osteoclasts (OCLs). For the training dataset, all TIFF-formatted images were auto-adjusted for brightness and contrast using ImageJ's default settings and then scaled down to reduce file size. Four representative images (σ = 10) were manually annotated using brush strokes to identify myeloid precursors (smaller, single magenta cells) and OCLs (larger, irregularly shaped, multinucleated magenta cells). Once the segmentation results accurately distinguished OCLs from precursors, the training file was used to develop a custom macro in ImageJ (version 1.54g) for automated measurement of total OCL area. Each segmented image was processed using minimum thresholding to separate foreground objects from the

background, followed by Gaussian blur ($\sigma = 2$) to smooth object edges. The images were then converted to binary format for particle analysis using ImageJ's 'Analyze Particles' function, applying a size threshold of 70 pixels to exclude small artifacts. Results (shown in Manuscript Fig. 6e) were expressed as the total area of OCLs (in square pixels) for each experimental condition.

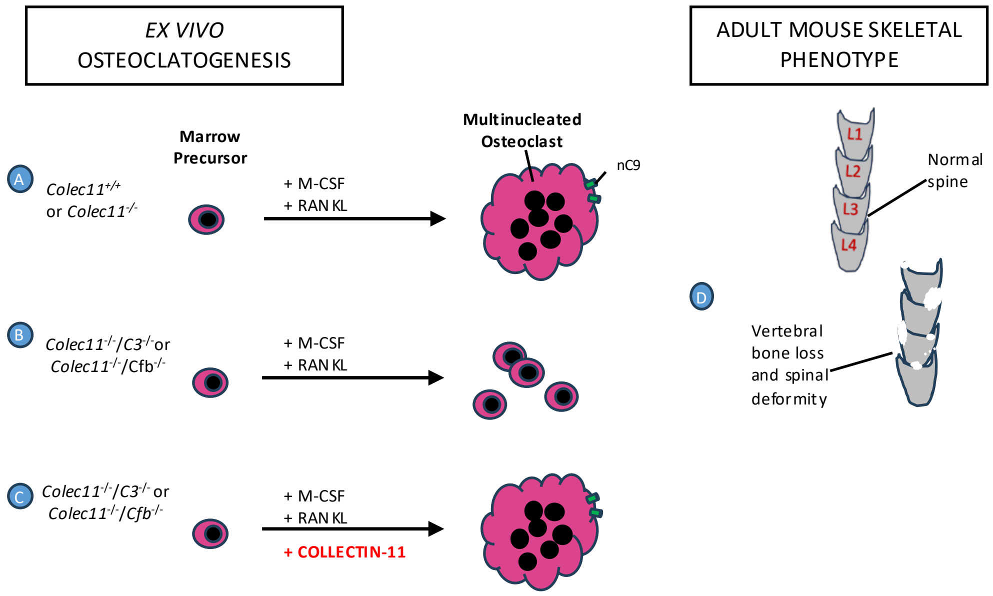


**Supplemental Figure 6. The Role of Collectin-11 in osteoclast development and bone maintenance.**
(A) Normal or CL-11-knockout bone marrow-derived stem cells (BMSC) cultured with M-CSF + RANKL differentiate into giant cell osteoclasts (OCL).
(B) Double knockout (DKO) mice lacking CL-11 *and* C3 *or* factor B exhibit failed OCL differentiation and MAC (nC9) formation.
(C) Supplementation with CL-11 restores OCL formation.
(D) DKO mice present with spinal deformity at 12 weeks old suggesting impaired bone maintenance.

### Tables

| Genotype | Total mice (15 litters) | Number of mice with kyphosis/scoliosis | % mice with kyphosis/scoliosis |
|---|---|---|---|
| WT | 101 | 0 | 0 |
| $C3^{-/-}$ | 79 | 0 | 0 |
| $Cfb^{-/-}$ | 81 | 0 | 0 |
| $Masp\text{-}2^{-/-}$ | 91 | 0 | 0 |
| $Colec11^{-/-}$ | 88 | 1 | 1.1 |
| $Colec11^{-/-}/C3^{-/-}$ | 78 | 22 | 28.2 |
| $Colec11^{-/-}/Cfb^{-/-}$ | 77 | 21 | 27.3 |
| $Colec11^{-/-}/Masp\text{-}2^{-/-}$ | 95 | 22 | 23.2 |
| $C3^{-/-}/Masp\text{-}2^{-/-}$ | 87 | 0 | 0 |

**Table 1. Frequency of kyphosis and scoliosis in wild-type (WT) and gene-deleted mice, as defined in the Methods section.**
Mice with double knockouts (DKO) that include the CL-11 deletion exhibit a higher rate of spinal abnormalities compared to single-knockout (SKO) mice and a DKO group lacking the CL-11 deletion (C3-/-/Masp-2-/-). Sample size: N = 77–101 mice per group, across 15 litters.

| Antibody | Source | Concentration |
|---|---|---|
| Rabbit anti human CL-11 | Abbexa (abx003772) | 1:100 |
| Mouse anti-human CL-11 | Provided by P. Garred | 1:100 |
| Rabbit anti-human C3d | Dako (A0063) (as previously (Howard et al., 2020, Farrar et al., 2016)) | 1:100 |
| Rabbit anti-rat C9 | Gift from P. Morgan | 1:100 |

**Table 2. List of antibodies used in the experiments, including their source and final concentration.**
